# Systemic sclerosis is a disease of a prematurely senescent, inflammatory and activated immunome

**DOI:** 10.1101/637736

**Authors:** Bhairav Paleja, Andrea Low Hsiu Ling, Pavanish Kumar, Suzan Saidin, Ahmad Lajam, Sharifah Nur Hazirah, Camillus Chua, Lai Li Yun, Salvatore Albani

## Abstract

Systemic sclerosis (SSc) is an autoimmune disease characterised by excessive fibrosis of skin and internal organs, and vascular dysfunction. Association of T and B cell subsets have been reported in SSc, however there is lack of systematic studies of functional relations between immune cell subsets in this disease. This lack of mechanistic knowledge hampers targeted intervention. In the current study we sought to determine differential immune cell composition and their interactions in peripheral blood of SSc patients. Mononuclear cells from blood of SSc patients and healthy controls (HC) were analysed by mass cytometry using a 36 marker (cell-surface and intracellular) panel. Transcriptome analysis (m-RNA sequencing) was performed on sorted T and B cell subsets. Unsupervised clustering analysis revealed significant differences in the frequencies of T and B cell subsets in patients. Correlation network analysis highlighted an overall dysregulated immune architecture coupled with domination of inflammatory senescent T cell modules in SSc patients. Transcriptome analysis of sorted immune cells revealed an activated phenotype of CD4 and MAIT cells in patients, accompanied with increased expression of inhibitory molecules, reminiscent of phenotype exhibited by functionally adapted, exhausted T cells in response to chronic stimulation. Overall this study provides an in-depth analysis of the systemic immunome in SSc, highlighting the potential pathogenic role of inflammation and chronic stimulation mediated “premature senescence” of immune cells.

**ONE SENTENCE SUMMARY:** The immune architecture in Systemic sclerosis is altered and reminiscent of chronic stimulatory environment led premature senescence of immune cells.

## INTRODUCTION

Systemic sclerosis (SSc) is a multi-systemic autoimmune disease defined by excessive microvascular damage, dysregulated immune system and increased deposition of extracellular matrix proteins leading to fibrosis of skin and internal organs (1). A lack of effective therapies is in part due to incomplete understanding of SSc pathogenesis, as well as inadequate mechanistic stratification of this heterogeneous disease. Traditional clinical subsetting according to the extent of skin thickening has shown that, in general, patients with limited cutaneous SSc (LcSSc) have a better prognosis than those with diffuse cutaneous SSc (DcSSc) (2). Although there are certain disease manifestations that differentiate these two groups, some of the severe organ manifestations, such as lung fibrosis, occur across groups, and are the cause of significant mortality.

Crosstalk between stromal cells like fibroblasts and immune cells is considered as a major mechanism of disease pathogenesis and progression. However the mechanisms responsible for the initiation of autoimmunity leading to fibrosis and the role of immune effector pathways in pathogenesis of SSc remain incompletely understood. Several studies have suggested the role of T and B cells in the pathogenesis of SSc. Detection of oligoclonal T cells in SSc suggest an antigen specific immune response (3) resulting from breakdown in self-tolerance and induction of T cell activation, secretion of pro-inflammatory and pro-fibrotic cytokines (4),(5). Indeed, increased numbers of T cells secreting pro-fibrotic and inflammatory cytokines have been detected in blood and affected skin of SSc patients. In addition, abnormalities in regulatory T cell are also implicated in pathogenesis of SSc (6). Apart from conventional CD4 and CD8 T cells, reports have also shown reduced numbers of “unconventional” mucosal associated invariant T (MAIT) and Gamma delta T cells in SSc patients, likely reflecting their mobilisation to inflamed skin (7),(8). Collectively, a dysfunctional immune system that is evident in the periphery seems to be a major component of SSc pathogenesis, lending to the hope of being able to distill clinically and mechanistically meaningful signatures from peripheral blood. However, most of the knowledge is fragmented, underscoring the need for an integrated understanding of the architecture of the Immunome in SSc, thus potentially leading to therapies targeting the aberrant immune responses (9).

The objective of this study is to build a map of the immunome in SSc, and to identify the perturbations which the disease causes by comparison with healthy controls. We employed for the purpose a combination of high dimensional technologies, namely CyTOF for the immune architecture and next generation RNA sequencing for the definition of the transcriptome in differential cell subsets identified by CyTOF. Our high dimensional approach revealed disease induced peculiarities in the immune architecture, with a polarisation towards activated, pro-inflammatory and senescent networks. More specifically, unsupervised clustering analysis revealed significant differences in the frequencies of T and B cell subsets in patients. Correlation network analysis highlighted an overall dysregulated immune architecture coupled with domination of inflammatory senescent T cell modules in SSc patients. Transcriptome analysis of sorted immune cells revealed an activated phenotype of CD4 and MAIT cells in patients, accompanied with increased expression of inhibitory molecules, reminiscent of phenotype exhibited by functionally adapted, exhausted T cells in response to chronic stimulation. Overall this study provides an in-depth analysis of the systemic immunome in SSc, highlighting the potential pathogenic role of inflammation and chronic stimulation mediated “premature senescence” of immune cells.

## RESULTS

### The architecture of the immunome is dysregulated in systemic sclerosis and pivots on inflammatory pathways

We exploited the high-dimensional, single cell analytical capability of CyTOF to define and dissect the immunome in SSc. Peripheral blood mononuclear cells (PBMC) from SSc patients and healthy controls were analysed using t-Distributed Stochastic Neighbour Embedding (tSNE) algorithm for dimension reduction, followed by clustering to identify nodes composed of similar cells. A comprehensive panel of 40 individual antibodies, specific for lineage, activation, cytokines, trafficking and differentiation, was designed and employed for the task (Supplementary Table 1). Figure 1A shows tSNE plot of the distribution of all the major immune lineages in SSc patients and HC. Comparing similar plots for patients and healthy controls (Figure 1B and 1C respectively) identified disease specific alterations in immune cell subsets in systemic sclerosis.

**Figure 1:**
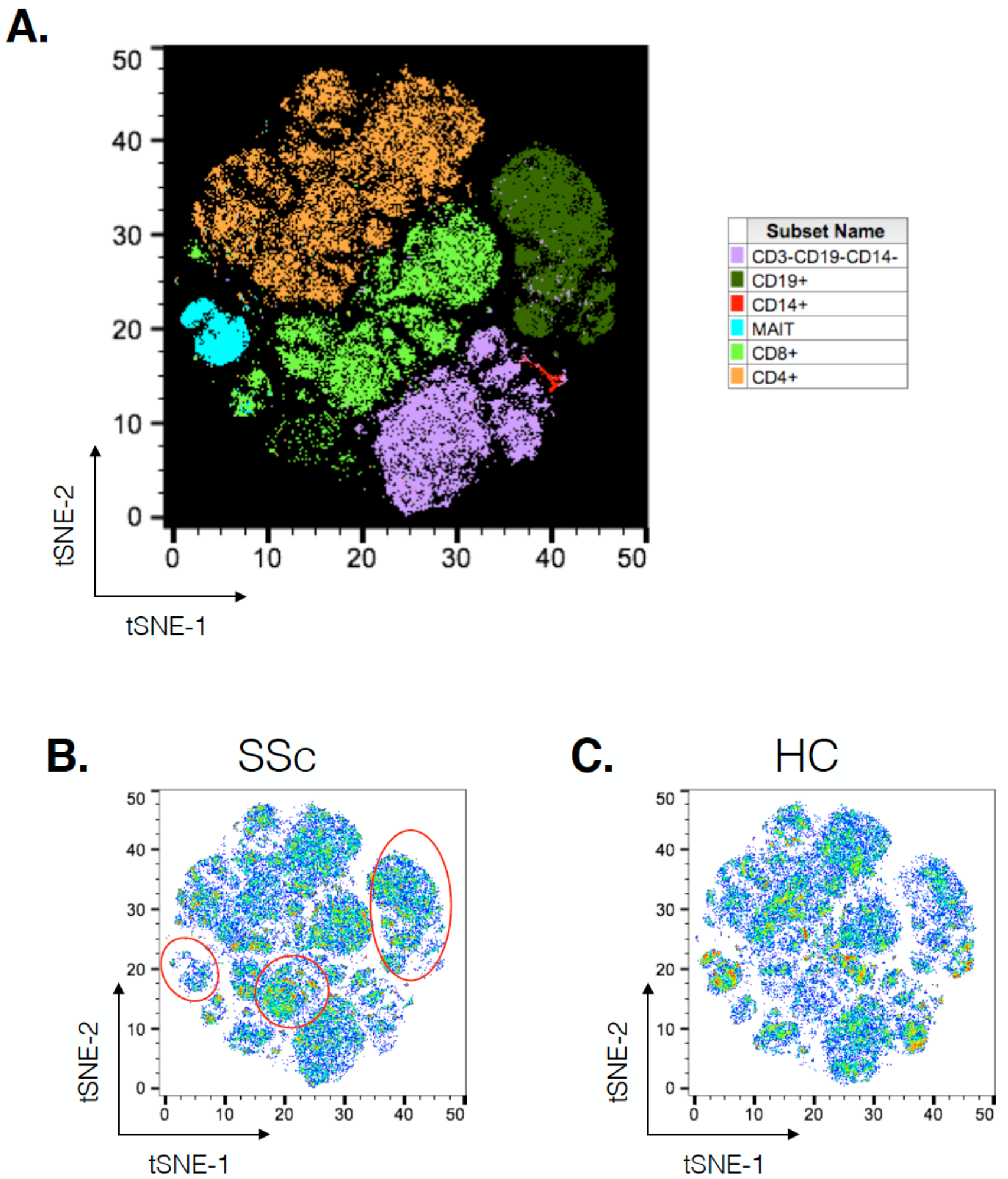
Unsupervised clustering reveals disease specific alterations in PBMCs from systemic sclerosis patients. Unsupervised clustering analysis of mass cytometry data from PBMCs of SSc patients (n=20) and healthy controls (n=10). Panel **A** tSNE maps for distribution of major immune subsets in SSc patients and healthy controls. All markers in CyTOF panel were used for clustering analysis. Panel **B** and **C** tSNE maps showing distribution of cells for SSc and HC respectively. Marked regions (red circle/ovals) highlight changes in cell subsets in patients vs healthy controls.

Unsupervised clustering analysis of CyTOF data from PBMCs of patients and HC identified a total of 133 nodes, with tight groupings of cells belonging to the same lineages. Comparison of frequencies of each node showed differences in immune subsets in SSc patients as compared to HC (Figure 2A). Statistical analysis revealed 18 nodes significantly different in SSc patients compared to HC (Supplementary Figure 1). This included nodes representing T cell (CD4+, CD8+ and MAIT) and B cell subsets (Figure 2B). This sort of analysis is very informative, but does not fully exploit the high dimensionality of CYTOF. Hence, we set off a relational analysis between different cell subsets, in order to define and understand if the architecture of the immunome in SSc could be a variation from the normal. To this end, we performed pairwise correlations between all nodes. Correlation values were used to define edges and create network between cellular nodes identified. Nodes were connected only if the absolute correlation values among them was >0.6. Network edges were colour coded to identify positive (green) and negative (red) correlations, with edge width determining absolute correlation. Network properties (Supplementary Table 2) were calculated and analysed to determine system level immune cell communication. Modularity score is a measure of network structure and reveals the strength of division of network in modules. Higher modularity score indicates a network with dense connections between nodes within modules but sparse connections between nodes in different modules. SSc patients showed higher modularity (0.26, Figure 2C) of nodes as compared to healthy controls (0.047, Figure 2D). Furthermore, the number of negatively correlated edges were reduced in patients (2.24%) as compared to healthy controls (30.4%). Another network property, centralisation score, is an indicator of dominance of nodes in overall network. Higher centralisation score was observed in patients cellular network (0.19) compared to HC network (0.031). To determine the identity of cell types in the different modules, we colour coded each node based on their phenotype. We found highly correlated distinct modules of B cells and NK like cells in SSc patients. In addition, tight modules of positively correlated CD4 and CD8 T cell subsets were evident in patients with such distinct clustering absent in immune networks from healthy controls. In summary, overall network analysis showed higher modularity, higher centralisation score, lower negatively correlated nodes and distinct cell type specific modules in cellular network from patients. These network properties are indicative of emergence of disease specific interactions in immune subsets and gearing of functions centralised around dominating nodes.

**Figure 2:**
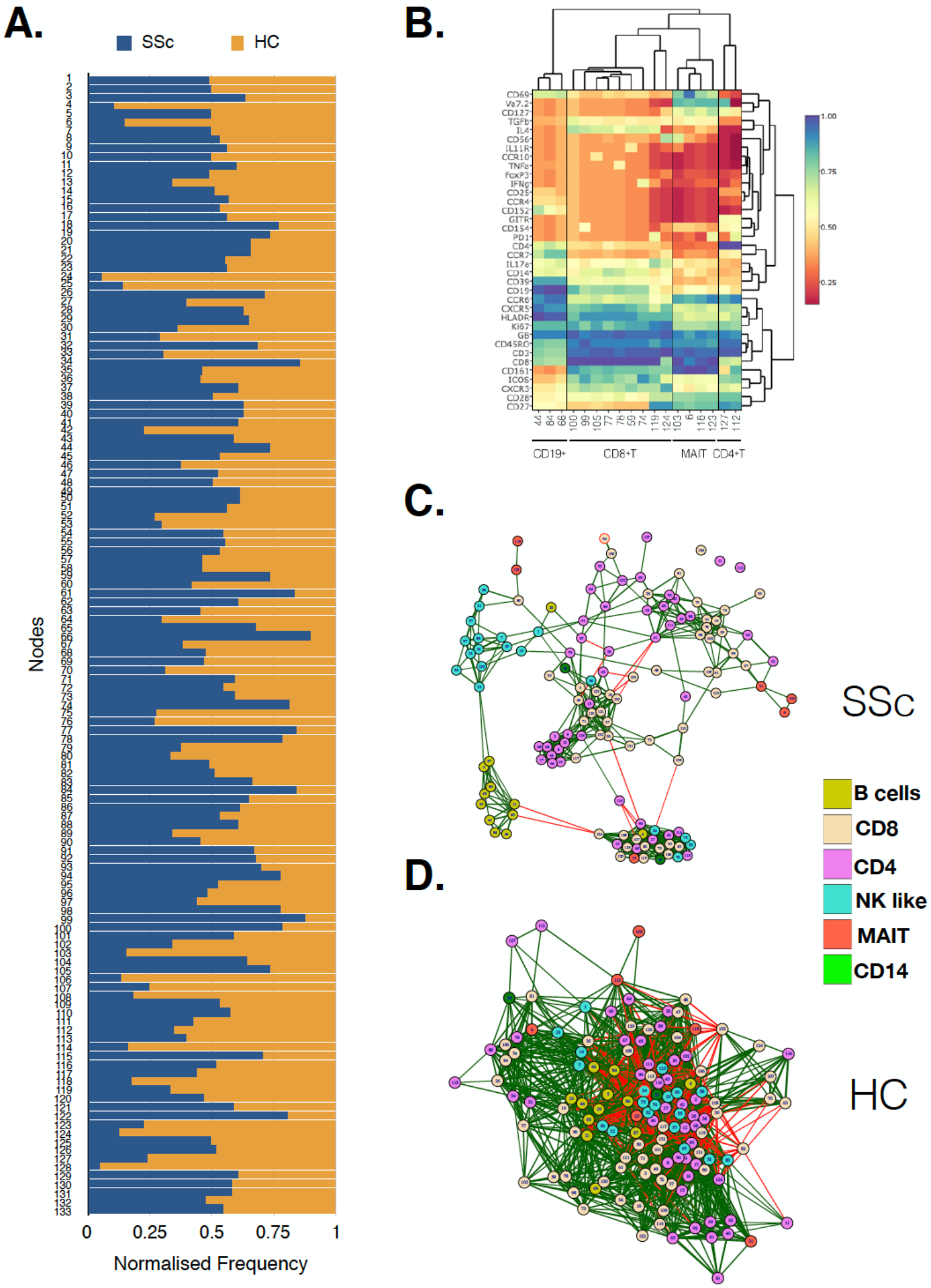
MAIT, CD4+ and CD8+ T cell subsets are affected in systemic sclerosis patients. Unsupervised clustering analysis of mass cytometry data from PBMCs of SSc patients (n=20) and healthy controls (n=10) was performed. Panel **A** Stacked bar chart shows normalised frequency of all nodes identified by unsupervised clustering analysis. **B**. Heatmap shows phenotype of nodes that were significantly (t-Test) different between SSc patients and HC. Median expression of all markers for each node was determined and plotted. **C.** Correlation network for all the nodes identified by unsupervised clustering in PBMCs from SSc patients. **D.** Correlation network for all the nodes identified by unsupervised clustering analysis in healthy controls. Colour of the nodes identify different cell types as indicated in legend. Green and red lines indicate positive and negative correlation, respectively.

As cellular network analysis of unstimulated immune cells from patients revealed differential modular structure and intercellular connections as compared to healthy controls, we further investigated cellular networks in patients and healthy controls based on their functional status. Each node from the mass cytometry data was classified based on the ability of cells in the node to produce cytokines. Thus the cells were identified as producing pro-inflammatory, anti-inflammatory, multiple cytokines, IL17 only or no cytokines. Cellular correlation network was then generated as previously described and nodes were color coded according to their functional responses. As observed for unstimulated cells, correlation network of stimulated cells from patients vs HC (Figure 3A) showed higher modularity (0.191 vs 0.037, respectively), lower negative correlations (0.73% vs 26.6%, respectively) and higher centralisation score (0.334 vs 0.039, respectively) as compared to healthy controls (Figure 3B, Table 3). Interestingly, cells producing IL17 formed a tight, highly correlated module in SSc patients but not in healthy controls. This, highly modular structure and grouping of IL17 producing cells in patients is suggestive of the prevalence of inflammatory functions in systemic sclerosis.

**Figure 3:**
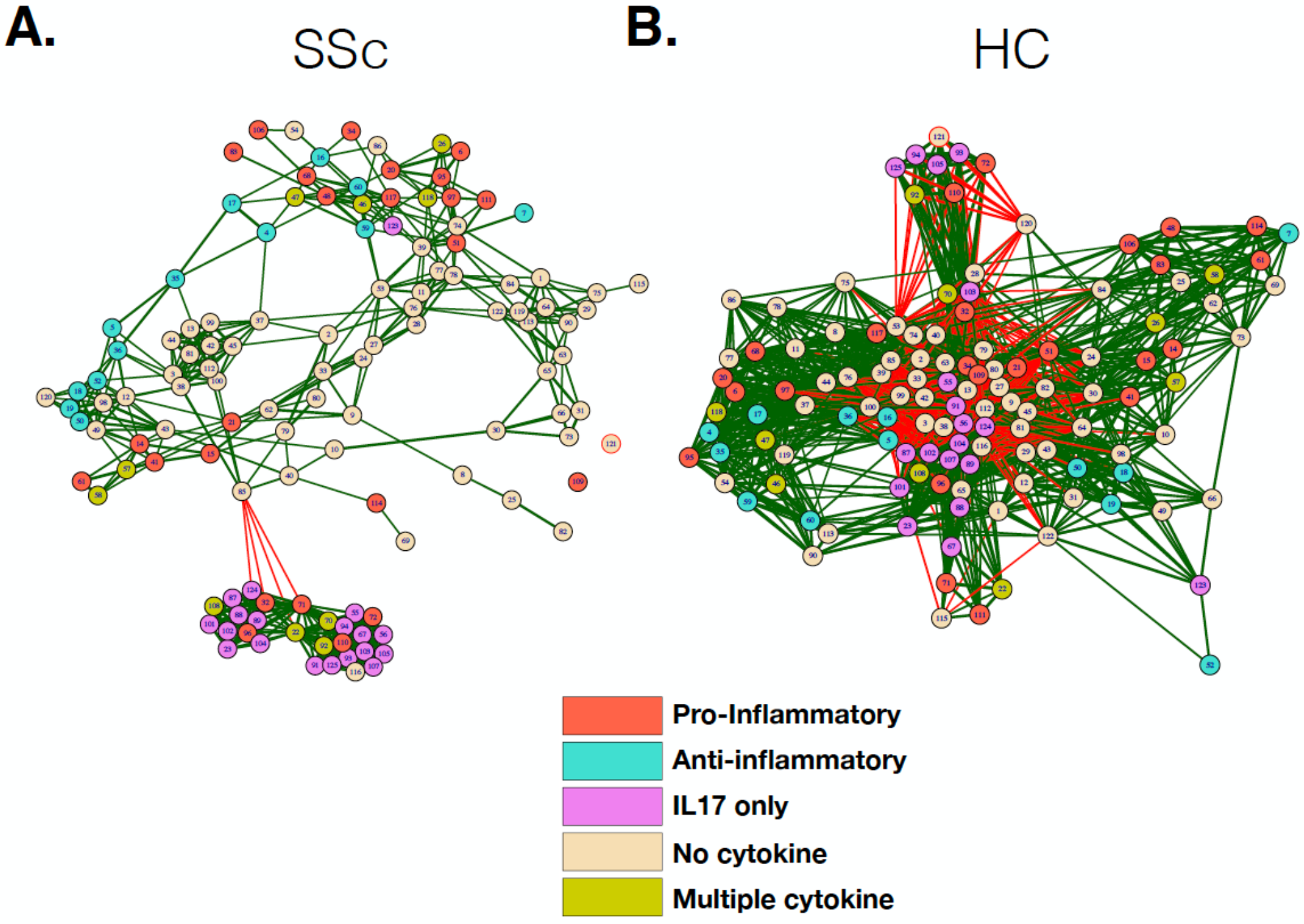
Correlation network shows enrichment of inflammatory cell subset modules in SSc patients. **A,B.** Correlation networks showing interaction between different cytokine producing cells from patients and healthy controls, respectively. Colour of the nodes identify cytokine producing capacity of cells in the particular node as indicated in legend. Green and red lines indicate positive and negative correlation, respectively. Anti- Anti-inflammatory, Pro-Pro-inflammatory, Multi- Multiple cytokines, NC- No cytokine, IL17 - IL17 producing cells.

### Gene signature reveals distinct functional state of T cells in systemic sclerosis, dominated by pro-inflammatory, activated, senescent cells

In order to better dissect the molecular mechanisms associated with the differences in the immune architecture in SSc patients, cell subsets which were differently represented in patients (i.e. CD4, CD8 T, MAIT and B cells, Figures 1 and 2) were sorted and subjected to transcriptome analysis by deep mRNA-sequencing. We performed principal component analysis to visualise overall transcriptome changes in sorted cells from SSc patients and HC. T cells from patients segregated distinctly from healthy controls (Figure 4A and 4B). Indeed, differential gene expression showed distinct gene expression profile in all the T cell subsets analysed, validating the finding by CyTOF that the immune mechanisms in SSc is functionally different than HC. In CD4 T cells, a total of 2266 genes were differentially regulated (1082 genes upregulated and 1184 genes down regulated) in SSc as compared to healthy controls (FDR cutoff <0.05 and minimum log2fold change = 2). In MAIT cells, total of 762 genes were differentially regulated, 549 unregulated and 213 down regulated in SSc as compared to healthy controls. In CD8 T cells, total of 20 genes were differentially regulated, 10 unregulated and 10 down regulated in SSc as compared to healthy controls. In B cells, at the set cutoff, there were no genes differentially expressed.

**Figure 4:**
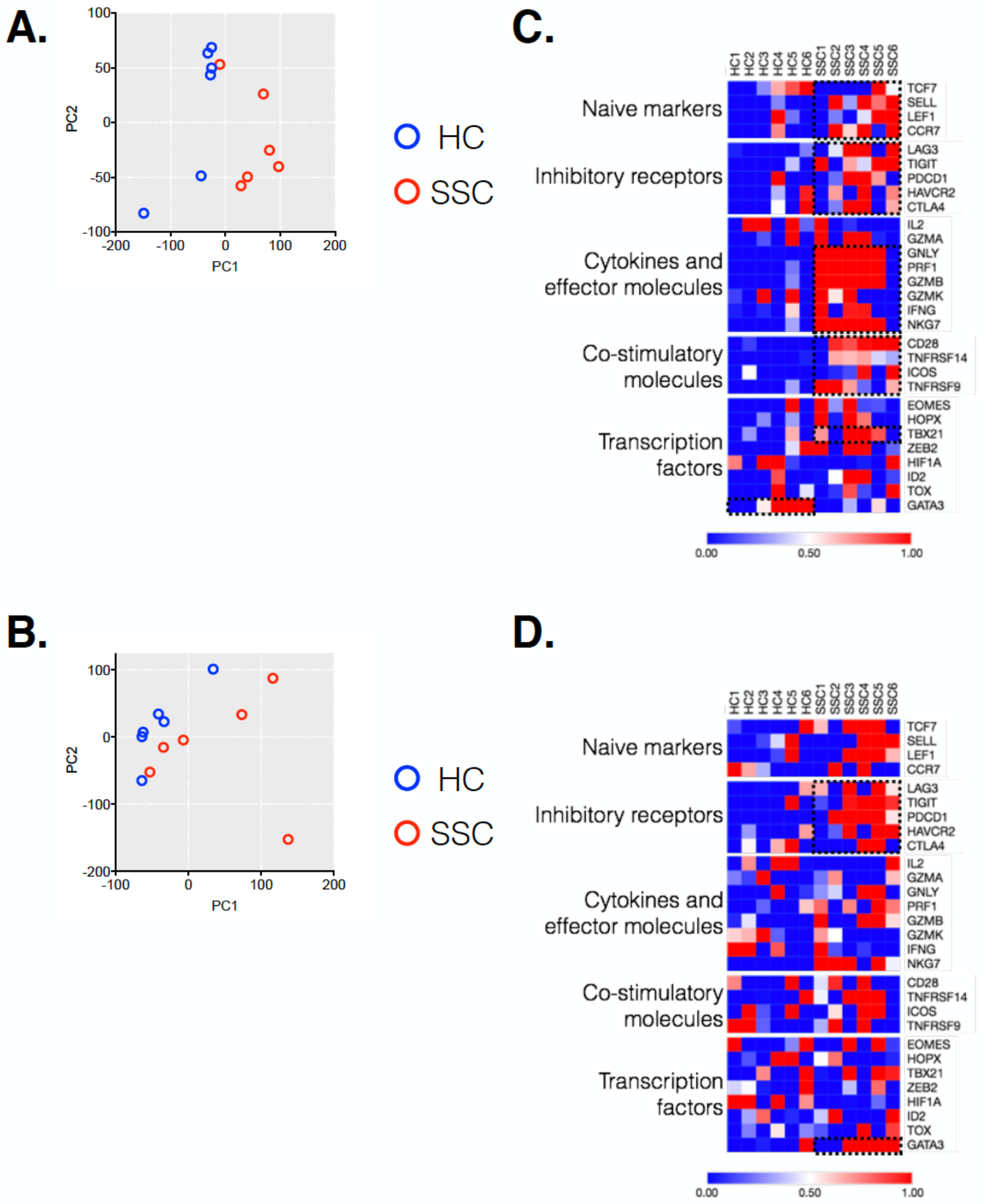
Transcriptome analysis of CD4 and MAIT cells reveals disease associated clustering of biological samples. **A, B.** Principal component analysis of CD4 T and MAIT cells, respectively, based on normalised read counts from RNA-seq data. Distinct separation of SSc patient samples and healthy controls is seen. **C, D.** Normalised median expression of selected T cell function associated genes in CD4 and MAIT cells, respectively.

Next, we delved in the mechanisms responsible for the differences in gene profiling. We analysed from the RNAseq dataset how differently expressed genes affect T cell activation and functional status (10). CD4 T cells from patients showed an increased expression of genes related to cytokines and effector molecules (Figure 4C). Interestingly, these samples also exhibited higher expression of inhibitory receptor genes such as PDCD1, TIGIT, LAG3 (Figure 4C). Similarly, MAIT cells from patients showed increased expression of inhibitory receptor genes (Figure 4D). The transcription factor GATA3 was highly expressed in MAIT cells from patients and not healthy controls. Conversely, CD4 T cells from patients showed increased expression of TBX21 as compared to healthy controls. Overall, transcriptome data revealed an activated phenotype of CD4 and MAIT cells in patients, though this was accompanied with increased expression of inhibitory/maturation molecules like PD1. Altogether, both the immunome and transcriptome profiles of T cell subsets eminent in SSc are reminiscent of “functionally adapted” senescent T cells in response to chronic stimulation, probably linking our findings to disease pathogenesis. We focused in particular on PD1 for its relevance in the cell responsiveness and homeostasis.

Indeed, in addition to its role in T cell exhaustion, PD1 has also been associated with expression on T cells under chronic stimulation (11). Moreover, elevated PD1 expression has been shown to be associated with attrition of MAIT cells in chronic infectious diseases (12). Our transcriptome data showed increased expression of PD1 on CD4 and MAIT cells from patients. To evaluate the role of increased PD1 expression in patients and its association with the frequency of various immune subsets in the periphery, we performed correlation analyses. PD1 expression on different T cell subsets (CD4+, CD8+, CD4−CD8− and MAIT cells) in our CyTOF data was determined by manual gating. Next we investigated the association between PD1 expression on T cell subsets with the frequency of these cells. Results showed that frequency of MAIT cells in periphery in patients was inversely correlated with PD1 expression on these cells (Figure 5A-C). Conversely, CD4− CD8− T cells were inversely correlated with PD1 expression in healthy controls but not in patients (Figure 5D).

**Figure 5:**
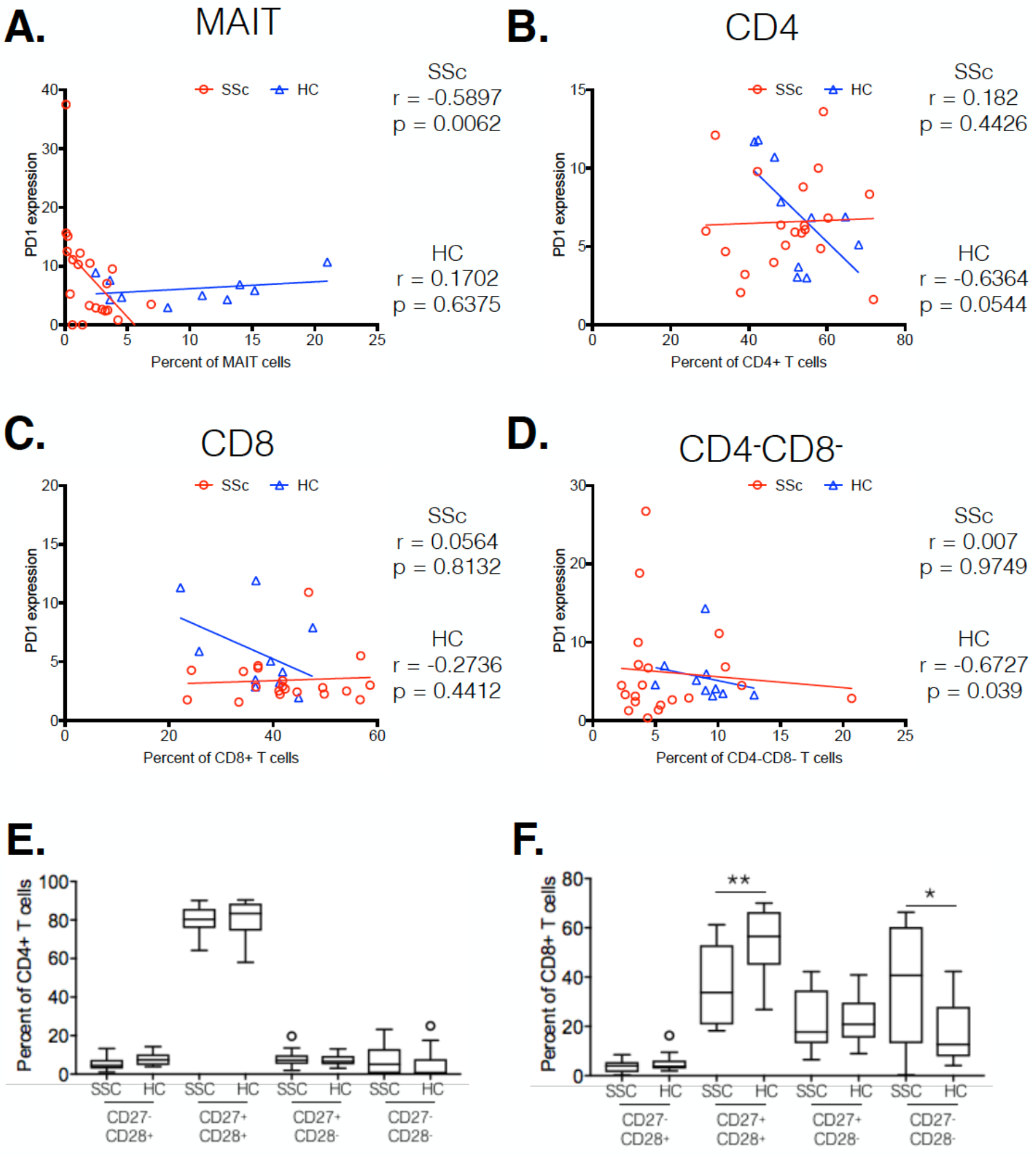
Systemic sclerosis is associated with PD1 expression and terminal differentiation of T cells. Correlation analysis between frequency of T cell subsets and respective PD-1 expression. Panel **A, B, C, D** shows MAIT cells, CD4+, CD8+ and CD4−CD8− T cells, respectively. Each point represents 1 sample. Correlation analysis was performed using Spearman’s rank correlation coefficient. **E, F.** Proportion of CD27/CD28 expressing cells in total CD4 and CD8 T cells from patients and healthy controls. (* p<0.05, ** p<0.01).

Taken together, transcriptome and CyTOF data pointed towards an immune architecture dominated by chronic antigen stimulated, senescent, pro-inflammatory rather than exhausted PD1+ T cells. To further ascertain this, we evaluated the differentiation state of CD4 and CD8 T cells. Based on the expression of co-stimulatory receptors CD27 and CD28, we observed an increased proportion of more differentiated CD27−CD28− cells in CD8 T cells from systemic sclerosis patients, with a concomitant decrease in CD27+CD28+ CD8 T cells (Figure 5E, F).

## DISCUSSION

This study was conceived to address the specific hypothesis that there are aberrations of the architecture of the peripheral immunome in SSc which are mechanistically relevant. We employed for the purpose a combination of high dimensional technologies and analysed the data in order to define spatially how the functional immune networks interlace.

Our approach departs from earlier efforts to understand the immune pathogenesis of systemic sclerosis, which have focused on individual subsets utilising conventional ways of analysis, providing a somewhat fragmented view of the immune landscape. There are contradictory reports on the frequencies of peripheral B cells in SSc patients. Indeed, the distribution of circulating B cells in SSc patients have shown to be increased (13), similar or decreased (14). The frequencies of B cell subsets (naive, memory and plasma blast) in our patients (Data not shown) confirmed earlier reports (13) showing increased naive B cells and decreased memory B cells in the periphery of systemic sclerosis patients. The decreased numbers of memory B cells have been attributed to an activated state and increased susceptibility to apoptosis (15).

MAIT cells represent a subset of unconventional T cell, characterised by invariant TCR repertoire and high expression of CD161. Since their identification recently, their role in infectious and autoimmune diseases is be becoming increasingly evident. Our observation of reduced MAIT cells in the periphery of SSc patients mirrors the finding by Mekinian et.al (8). Similar findings have been reported in other conditions including autoimmune diseases like inflammatory bowel disease, Sjogren’ syndrome, ankylosing spondylitis (AS), rheumatoid arthritis (RA), ulcerative colitis and SLE ((16);(17);(18);(19);(20)) and infectious diseases like HIV and HCV ((21); (22)). Gracey et. al. (18) reported, a decreased MAIT cell frequency in AS and RA patients that likely reflected recruitment to site of inflammation such as gut or inflamed joint.

Newer high dimensional technologies like mass cytometry and NGS makes it possible to produce highly correlated data sets that capture the modularity and inter cellular dynamics of immune system. Such a holistic approach enabled the identification, at the systems-level rather than mono dimensionally, of the most relevant immune components responsible for pathogenesis of disease.

Our correlation network analysis revealed segregation of immune cell subsets into modules of correlated immune features and functions. We found a higher modularity, lower negatively correlated nodes, higher centralisation score and distinct cell type specific modules in SSc immune cell network compared to healthy controls, implying disease specific interactions between immune subsets. Functional data also highlighted increased interplay between IL17 producing cellular subsets in SSc patients compared to healthy controls. Our data are validated by reports that IL17 and IL17 producing cells are increased not only in the periphery but also the skin in SSc ((23); (24); (25); (26)). Yang et.al. (27) showed also induction of fibroblast growth and increased collagen expression and protein secretion by IL17. Our approach brings these observations into a functional and possibly pathogenic framework. Although further studies evaluating the role of IL17 and IL17 producing cells in SSc are warranted, overall, our data show that highly correlated immune cell subsets and functional modules are pivoting on IL17 with obvious potential pathogenic and therapeutic implications.

Our approach also opened a new perspective on the immune pathogenies of the disease. In this study we observed striking changes in transcripts of CD4 T and MAIT cells from SSc patients. Targeted analysis using a set of T cell specific functional genes revealed increased expression of inhibitory-senescence receptors in CD4 and MAIT cells from SSc patients. Expression of co-inhibitory receptors, such as PD-1, LAG3, TIM-3 and TIGIT, on immune cells is crucial for the regulation of peripheral tolerance (28). Studies have reported increased expression of inhibitory receptors on immune cell subsets in various autoimmune diseases including SLE ((29); (30)), psoriatic arthritis (31) and rheumatoid arthritis (32). Fleury et. al. (33) demonstrated increased expression and altered activity of inhibitory receptors in lymphocytes from SSc patients. Our transcriptome data highlighted changes in expression of such receptors on CD4 and MAIT cells in our patients. In addition, correlation analysis of mass cytometry data, revealed an inverse correlation between PD1 expression and frequency of MAIT cells in SSc patients compared to healthy controls. In interpreting these data in a functional perspective, it must be underscored that expression of inhibitory receptors has been linked to reduced cytokine production by T cells in viral infections and cancer patients ((34), (35)). However, many recent studies have also shown that the expression of receptors like PD-1 does not correlate with lower functionality but rather differentiation state. True to this, our mass cytometry data showed an increased frequency of terminally differentiated subsets of CD27−CD28− T cells in patients. The combination of our high dimensionality data identified cell subsets reminiscent of functionally adapted, exhausted T cells in response to chronic stimulation.

Altogether, the combination of these data points toward a dominance in SSc of T cell subsets which are highly differentiated, chronically stimulated and as such senescent, yet functional. Overall this study delineates for the first time the architecture of systemic immunome in SSc, highlighting the roles of inflammation and chronic stimulation-mediated “premature senescence” of immune cells. The knowledge distilled from this study has the dual translational valency of providing novel tools for monitoring and manipulating the deranged Immunome in SSc.

## MATERIALS AND METHODS

### Patients

Blood samples were collected from 23 patients with SSc (from the Scleroderma Clinic, Department of Rheumatology and Immunology at Singapore General Hospital) and 10 healthy controls (HC). Patients fulfilling the 2013 American College of Rheumatology/European League Against Rheumatism criteria were included in this study. Of 23 patients enrolled, 21 patients (91%) were female. The mean age at disease onset from first non-Raynaud’s phenomenon symptom was 41 years. Mean disease duration at time of blood collection was 5.2 years. Approximately half of the patients had LcSSc (n=12) or DcSSc (n=11). The frequency of anti-centromere antibodies and anti-topoisomerase-I antibodies was 3 and 12, respectively. The patients’ clinical characteristics are summarised in Table 1. This study was approved by Institutional review board of Singapore General hospital. All patients signed informed consent to participate in the study.

**Table 1:** Patient clinical characteristics

| Patient No. | Age | Gender | SSc subtype | Autoantibodies | Disease duration (Years) | Medication | Organ involvement |
| --- | --- | --- | --- | --- | --- | --- | --- |
| 1 | 59 | F | LcSSc | ACA | 11.70 | No | V, GI |
| 2 | 48 | M | LcSSc | Topo-I | 3.13 | P, MMF | V, GI, ILD |
| 3 | 50 | F | LcSSc | Topo-I | 5.21 | CYC | M, V, GI, ILD |
| 4 | 42 | F | LcSSc | No | 1.37 | No | M, V, GI |
| 5 | 42 | F | LcSSc | ACA | 1.52 | P | M, V, GI |
| 6 | 35 | M | LcSSc | Topo-I | 0.44 | P | Cardiac, M, V, GI, ILD |
| 7 | 60 | F | DcSSc | Topo-I | 1.16 | No | M, V, GI, ILD |
| 8 | 61 | F | DcSSc | No | 0.68 | No | V, GI, ILD |
| 9 | 24 | F | DcSSc | No | 3.26 | No | Cardiac, M, V, GI |
| 10 | 45 | F | DcSSc | No | 0.27 | No | GI |
| 11 | 52 | F | DcSSc | No | 3.45 | MTX | M, V, GI |
| 12 | 31 | F | LcSSc | Topo-I | 0.96 | P, MMF | M, GI, ILD |
| 13 | 63 | F | LcSSc | Topo-I | 6.84 | No | M, V, GI |
| 14 | 39 | F | DcSSc | Topo-I | 11.79 | CYC | M, V, GI |
| 15 | 42 | F | DcSSc | Topo-I | 2.24 | P, CYC | Cardiac, M, V, GI, ILD |
| 16 | 60 | F | DcSSc | Topo-I | 9.56 | MMF | M, V, ILD |
| 17 | 22 | F | DcSSc | Topo-I | 1.00 | No | M, V, GI, ILD |
| 18 | 47 | F | DcSSc | No | 3.91 | P, MMF | M, V, GI, ILD |
| 19 | 62 | F | LcSSc | No | 10.93 | No | M, V, GI, ILD |
| 20 | 55 | F | LcSSc | No | 26.53 | No | M, V, GI |
| 21 | 33 | F | LcSSc | ACA | 2.43 | P, MMF | M, V, GI |
| 22 | 59 | F | LcSSc | Topo-I | 10.45 | MTX | M, V, GI, ILD |
| 23 | 36 | F | DcSSc | Topo-I | 0.27 | MMF | M, GI |

\*Disease duration defined from first onset of non-Raynaud's phenomenon symptom; F - female; M – male; LcSSc - limited cutaneous SSc; DcSSc - diffuse cutaneous SSc; ACA - Anti-centromere antibody; Topo-I - Anti-topoisomerase I antibody; P- Prednisolone; MMF - Mycophenolate mofetil; MTX - Methotrexate; CYC - Cyclophosphamide; GI - Gastrointestinal; ILD - Interstitial lung disease; M - Musculoskeletal; V- Vasculopathy

### Isolation of Peripheral blood mononuclear cells

Peripheral blood was collected from all subjects in K2EDTA tubes and PBMCs were isolated by density gradient centrifugation using Ficoll-Paque Plus (GE Healthcare). Cells were resuspended in freezing medium containing 90% FBS and 10% DMSO, aliquoted and stored in liquid nitrogen until use.

### Mass cytometry staining and analysis

PBMCs were analysed with 40 metal conjugated antibodies specific for cell surface and intracellular antigens using CyTOF. Samples were thawed and left un-stimulated or were stimulated with phorbol 12-myristate 13-acetate (PMA) and Ionomycin for 4-hours. Brefeldin A and Monensin was added to culture in the last 3 hours of incubation. The cells were stained with cisplatin to identify live cells. Next, a combination of four anti-CD45 antibodies were used as barcodes (36) to allow simultaneous sample processing and reduce technical variability. This was followed by incubation with metal conjugated surface antibodies, fixation with 1.6% paraformaldehyde, permeabilization with 100% methanol and intracellular metal conjugated antibodies. Finally the cells were labeled with iridium containing DNA-intercalator. Samples were acquired on CyTOF 2 mass cytometer (Fluidigm).

EQ Four element calibration beads were used for signal normalisation. Before downstream analysis live cells were gated manually on event_length, ^195^Pt and DNA (^191^Ir and ^193^Ir) and debarcoded using FlowJo software (Supp. Figure 2). The debarcoded files were down-sampled to 5000 cells each. These files were analysed using an in-house developed software based on Barnes-Hut SNE non linear dimension reduction algorithm followed by k-means clustering (37). Heatmaps to identify phenotypes of significantly different nodes were plotted in R programming environment.

### Next generation sequencing

Total RNA was isolated using a Picopure RNA-Isolation kit (Arcuturus, Ambion) and cDNA was generated using SMART Seq® v4 Ultra™ Low Input RNA Kit (Clontech), both according to manufacturer’s instructions. Illumina-ready libraries were prepared from cDNA using the Illumina Nextera XT DNA Library Prep Kit (Illumina, SD). Next generation sequencing (NGS) was performed at the NGS Platform of the Genome Institute of Singapore (GIS) using 2×151bp on a HiSeq 4000 platform.

### RNA-sequencing data processing

Raw RNA-sequencing reads were mapped to the human genome (Hg19) using STAR aligner (38) with default parameter. The counts of reads mapped over gene features were obtained using the featureCounts method of the Subread package (39). Fold change was calculated in the R statistical programming software. exactTest built in the edgeR package and recommended for two group test was used for differential gene expression analysis.

### Network analysis

Correlation Network analysis was performed as previously described (40). Briefly, proportion of each node was calculated for every sample and the correlation between nodes was determined for SSc and HC. To construct the network, nodes were connected if they had an absolute correlation coefficient >0.6. The network was visualised and analysed using the igraph R package.

### Statistical Analysis

For CyTOF data, non-parametric unpaired Mann Whitney tests (GraphPad Prism) were used to identify differential nodes between the 2 groups. A p<0.05 was considered statistically significant.

## SUPPLEMENTARY MATERIALS

Supplementary Figure 1: Gating strategy employed

Supplementary Figure 2: Differentially expressed nodes in systemic sclerosis

Supplementary Table 1: CyTOF staining panel

Supplementary Table 2: Network properties

## Acknowledgements

The authors would like to thanks Dr. Lakshmi Ramakrishna for contribution in scientific discussion and comments on manuscript.

## Funding

This study was supported by grants to S.A from the NMRC (NMRC/STaR/020/2013, NMRC/MOHIAFCAT2/2/08, MOHIAFCAT 2 / 0001 / 2014, Centre Grants, TCR 15 Jun 006, NMRC/ CIRG/ 1460 / 2016, MH 095:003\016-0002), Duke-NUS, A*STAR BMRC (IAF311020), BMRC (SPF2014/005), Duke-NUS and SingHealth core funding. This research was also supported by the National Research Foundation Singapore under its NMRC Centre Grant Program (NMRC/CG/M003/2017) and administered by the Singapore Ministry of Health’s National Medical Research Council. Also supported by NMRC grant NMRC/CIRG/1426/2015 to A.L.H.L

## Author contributions

B.P performed the experiments, analysed the data and wrote the manuscript. P.K. performed bioinformatics and data analysis. S.S, A.L, S.N.H, C.C, L.L.Y, performed the experiments. A.L.H.L recruited the patients and obtained the relevant blood samples. S.A and A.L.H.L conceived the study, analysed the data and wrote the manuscript.

## Conflicts of interest disclosure

The authors declare no conflicts of interest.

