## Supplementary materials for "Systemic sclerosis is a disease of a prematurely senescent, inflammatory and activated immunome"

**Supplementary Figure 1:** Differentially expressed nodes in systemic sclerosis patients

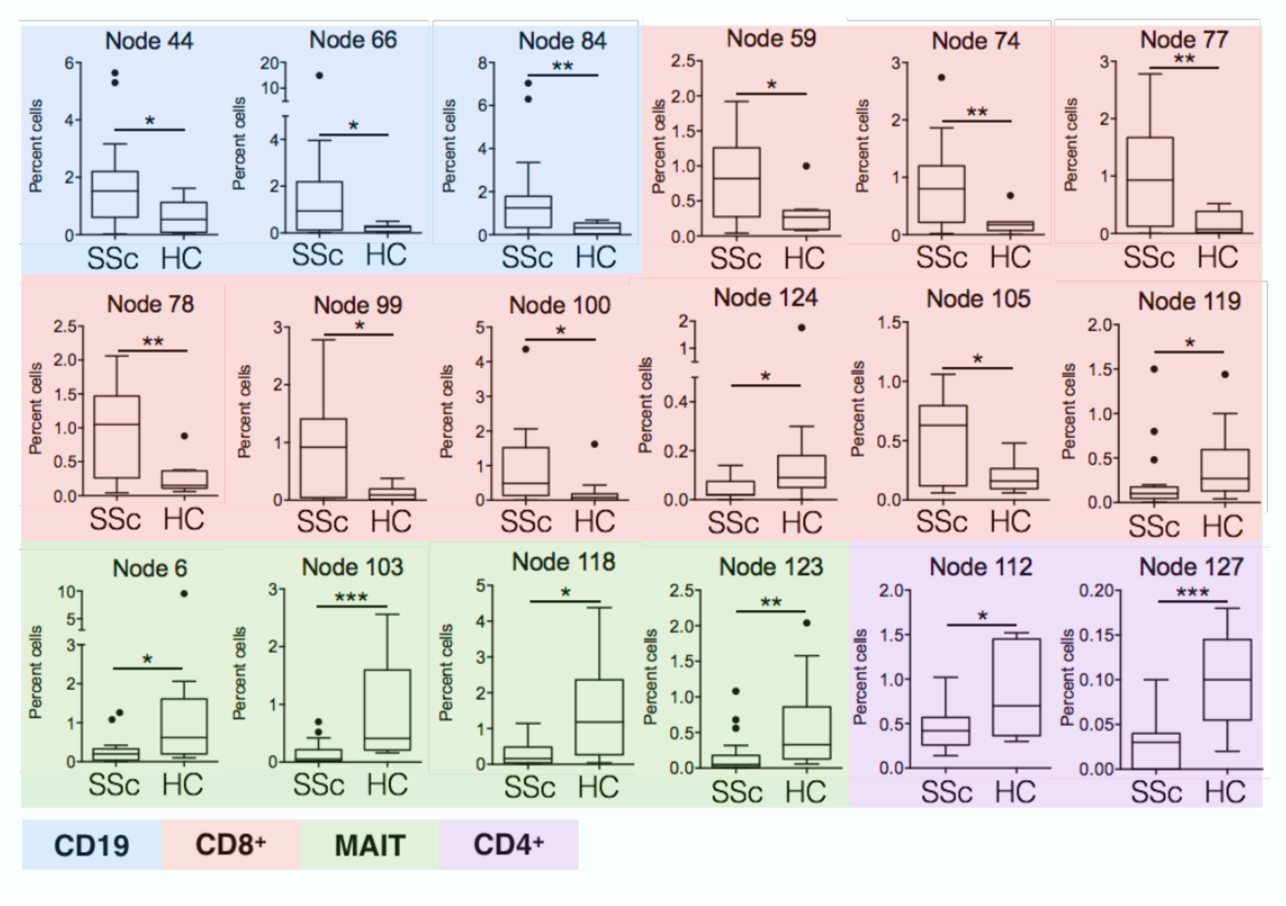


**Supplementary Figure 2 :** Gating strategy employed

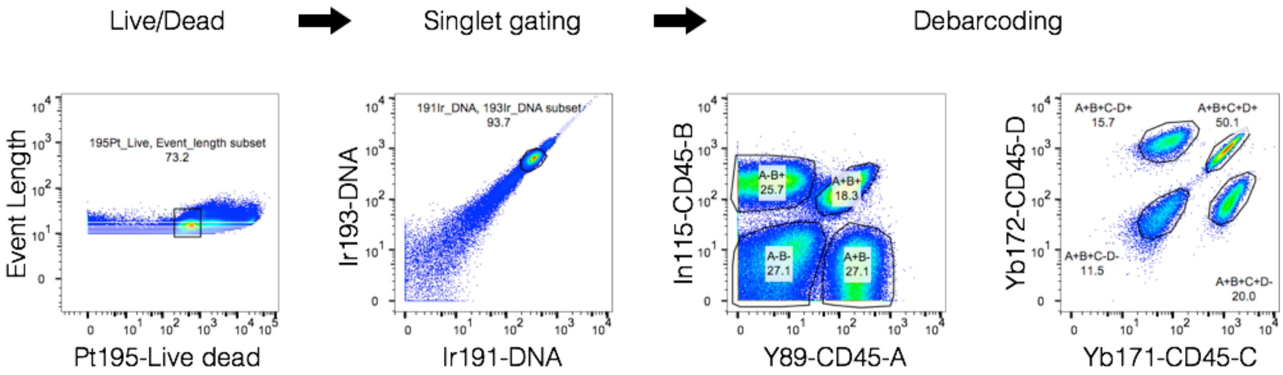

**Supplementary Table 1:** CyTOF staining panel

| Metal | Antibody | Clone | Vendor |
|---|---|---|---|
| **89** | CD45A | HI30 | Fluidigm |
| **112/114** | CD14 | TuK4 | Lifetechnologies |
| **115** | CD45B | HI30 | Biolegend |
| **139** | HLA-DR | L-243 | Biolegend |
| **141** | CD39 | A1 | Biolegend |
| **142** | CD45RO | UCHL1 | Biolegend |
| **143** | CD3 | UCHT1 | Biolegend |
| **144** | CD8 | SK1 | Biolegend |
| **145** | IL-4 | 8D4-8 | Biolegend |
| **146** | CD28 | CD28.2 | Biolegend |
| **147** | PD-1 | EH12.2H7 | Biolegend |
| **148** | CD4 | SK3 | Biolegend |
| **149** | CCR10 | 6588-5 | Biolegend |
| **150** | CD25 | 2A3 | Biolegend |
| **151** | CD56 | HCD56 | Biolegend |
| **152** | TNF-a | Mab11 | Biolegend |
| **153** | TGF-b | TW4-2F8 | Biolegend |
| **154** | CD27 | O323 | Biolegend |
| **155** | CD152 | BN13 | BD Pharmingen |
| **156** | CD127 | A019D5 | Biolegend |
| **157** | CCR4 | L291H4 | Biolegend |
| **158** | CD154 | 24-31 | Biolegend |
| **159** | CXCR5 | J252D4 | Biolegend |
| **160** | CD161 | HP-3G10 | Biolegend |
| **161** | CCR7 | G043H7 | Biolegend |
| **162** | FoxP3 | PCH101 | eBioscience |
| **163** | CXCR3 | G025H7 | Biolegend |
| **164** | GITR | 621 | Biolegend |
| **165** | IL11R | EPR5446 | Biolegend |
| **166** | Ki67 | 20Raj1 | eBioscience |
| **167** | ICOS | 2D3 | Biolegend |
| **168** | IFN-y | B27 | Biolegend |
| **169** | IL-17A | BL168 | Biolegend |
| **170** | CCR6 | G034E3 | Biolegend |
| **171** | CD45C | HI30 | Biolegend |

| Metal | Antibody | Clone | Vendor |
|---|---|---|---|
| **172** | CD45D | HI30 | Biolegend |
| **173** | GranzymeB | CLB-GB11 | Abcam |
| **174** | CD19 | HIB19 | Biolegend |
| **175** | Va7.2 | 3C10 | Biolegend |
| **176** | CD69 | FN50 | Biolegend |
| **191/193** | DNA | | |
| **195** | Live/Dead | | |

**Supplementary Table 2 :** Network properties

| Network property | Non stimulated | | Stimulated | |
|---|---|---|---|---|
| | SSc | HC | SSc | HC |
| **Modularity Score** | 0.26 | 0.047 | 0.191 | 0.037 |
| **Network centralization** | 0.19 | 0.031 | 0.334 | 0.039 |
| **Network density** | 0.065 | 0.24 | 0.070 | 0.232 |
| **Avg. no. of neighbours** | 9.2 | 34.59 | 8.72 | 28.8 |
| **Negatively correlated edges** | 13 | 659 | 4 | 480 |
| **Positively correlated edges** | 566 | 1503 | 541 | 1320 |
| **% negatively correlated edges** | 2.24 | 30.4 | 0.73 | 26.6 |
